# Metage2Metabo: metabolic complementarity applied to genomes of large-scale microbiotas for the identification of keystone species

**DOI:** 10.1101/803056

**Authors:** Arnaud Belcour, Clémence Frioux, Méziane Aite, Anthony Bretaudeau, Anne Siegel

## Abstract

Capturing the functional diversity of microbiotas entails identifying metabolic functions and species of interest within hundreds or thousands. Starting from genomes, a way to functionally analyse genetic information is to build metabolic networks. Yet, no method enables a functional screening of such a large number of metabolic networks nor the identification of critical species with respect to metabolic cooperation.

Metage2Metabo (M2M) addresses scalability issues raised by metagenomics datasets to identify keystone, essential and alternative symbionts in large microbiotas communities with respect to individual metabolism and collective metabolic complementarity. Genome-scale metabolic networks for the community can be either provided by the user or very efficiently reconstructed from a large family of genomes thanks to a multi-processing solution to run the Pathway Tools software. The pipeline was applied to 1,520 genomes from the gut microbiota and 913 metagenome-assembled genomes of the rumen microbiota. Reconstruction of metabolic networks and subsequent metabolic analyses were performed in a reasonable time.

M2M identifies keystone, essential and alternative organisms by reducing the complexity of a large-scale microbiota into minimal communities with equivalent properties, suitable for further analyses.

## Background

Understanding the interactions within microbiotas is crucial for ecological [53] and health [11] applications. With the improvements of metagenomics, and in particular the rise of methods to assemble individual genomes from metagenomes, unprecedented amounts of data are available to disentangle the functioning of microbiotas. This provides ways to tackle the potential role of microbes, whereas previous metataxonomics [34] analyses could only provide information on “who is there”. Henceforth, the main challenge is to handle both the scale of metagenomics datasets, and the incompleteness of their data. Hundreds or thousands of genomes can be reconstructed from various environments [41, 16, 61, 52], either with the help of reference genomes or through metagenome-assembled genomes (MAGs).

These genomes are the starting point to a large set of analyses dedicated to gather the functions of the considered organisms, and possibly study them with regard to functions performed by a host. A first level of analyses is to annotate these genomes and characterise the families of molecular processes likely to happen in the species, which can rely on ontologies [4, 24]. Another possibility is to target the whole metabolism and build a genome-scale metabolic network (GSMN) for each individual genome.

GSMNs gather all the expected metabolic reactions of an organism. Thiele et al [54] defined a precise protocol for building them, associating the use of automatic methods and a thorough curation of the model, based on expertise, literature, and mathematical analyses. This has been the basis for many implementations of GSMN reconstruction as all-in-one platforms [13, 27, 3], toolboxes [1, 57, 46], or individual tools for targeted refinements and analyses on GSMN [43, 55, 56]. Automatic reconstruction of GSMNs relies on the annotation of the genomes, as well as on the search for orthologues in known species. As expected based on the initial protocol for reconstruction described in [54], and despite the improvements of methods since then, automatic reconstructions of GSMN are likely to be incomplete. They produce GSMN drafts that need to be further refined and/or gap-filled, requiring human expertise that cannot be easily contemplated for a large number of genomes. [28]. Nevertheless, in the context of microbiotas and poorly-described organisms, it is hypothesised that some gaps in the metabolism can be explained by the dependency of organisms to each others [36]. It is therefore relevant to systematically analyse the complementarity between metabolic networks of microbiota species as a index for putative cooperation between them [58].

As metagenomics generates a large number of genomes for bacteria having the capability of exchanging compounds, their interactions deserve to be investigated at the metabolic scale. It is a challenging objective that entails turning hundreds or thousands of genomes into metabolic information, and analysing the latter according to their capabilities of individually or collectively produce metabolic compounds. Such a metabolic screening of organisms capabilities in microbiotas should enable the global comprehension of the functions expected in each member, based on its genetic content. Notice however that the objective of considering all the species together and study the complementarity of their respective metabolisms advocates for a very restrictive use of gap-filling procedures of individual metabolic networks. They indeed may artefactually assume that individual organisms each sustain their own growth in a restricted environment, whereas they rely on metabolic interactions to do so. As GSMNs are built individually, the purpose is instead to perform metabolic analyses on genomes that have been homogeneously prepared and take the most out of automatically built draft GSMNs, in order to highlight differences between them.

A variety of toolboxes have been proposed to study communities of organisms with GSMNs [30, 48]. Some studies are focused on pairwise interactions. Analyses on larger communities mainly rely on constraint-based modeling [8, 60, 29] at steady or unsteady states. However, these tools were applied to small size communities, usually no more than ten members, which suggests a computational bottleneck that needs to be faced for larger communities [30]. In addition, the GSMNs have to be of high-quality for accurate mathematical predictions. Therefore, as stated by [30], development of tools tailored to the analysis of large communities is needed. Being able to identify the main metabolic features of organisms beyond functional annotation is critical in microbiotas for the identification of important species, and further experiments or curation of their models.

Here we describe a software, metage2metabo (M2M), to analyse metabolic complementarity starting from individual annotated genomes in large microbiotas. M2M first characterises the added-value of organisms complementarity in terms of metabolic compounds production. Then it identifies communities and keystone species with respect to a family of metabolic compounds selected from the previous step. M2M uses the algorithm of network expansion [14] to capture the set of producible metabolites in a GSMN, therefore handling stoichiometry inaccuracy which is commonly faced with automatically reconstructed models. We therefore advocate for the identification of keystone species and the added-value of cooperation within the microbiota using M2M to evaluate the individual GSMNs and screen the collective metabolism.

To illustrate the metabolic screening potential of our method, we selected two large-scale sets of genomes for analysis: 913 cow rumen MAGs [52], and a set of 1,520 draft bacterial reference genomes from the gut microbiota [61]. We show that M2M can efficiently reconstruct metabolic networks for each genome, identify potential metabolites produced by cooperating bacteria, and suggest minimal communities and keystone species associated to their production.

## Implementation

Metage2Metabo (M2M) is a Python package. It can be used on a personal computer or on a cluster (using the Python package, Docker or Singularity) to benefit from its multi-processing functionality on large microbiotas datasets. A detailed documentation is available on metage2metabo.readthedocs.io.

M2M’s main pipeline consists in three main steps performed sequentially: i) automatic reconstruction of metabolic networks for a large number of annotated genomes, ii) analysis of metabolic capabilities for each metabolic network and computation of the cooperation potential, i.e. the set of metabolites predicted to become producible through complementarity of synthetic pathways, and iii) identification of minimal communities and keystone species for a targeted set of compounds.

Figure 1 depicts the pipeline of M2M. The inputs for the whole workflow are a set of annotated genomes, and a list of nutrients representing a growth medium. However, each step can also be run individually. For instance, one can perform the metabolic network analysis by providing the GSMNs and the growth medium as inputs, or the community selection by providing the GSMNs, the medium and a family of metabolic compounds aimed to be produced by the community.

**Figure 1:**
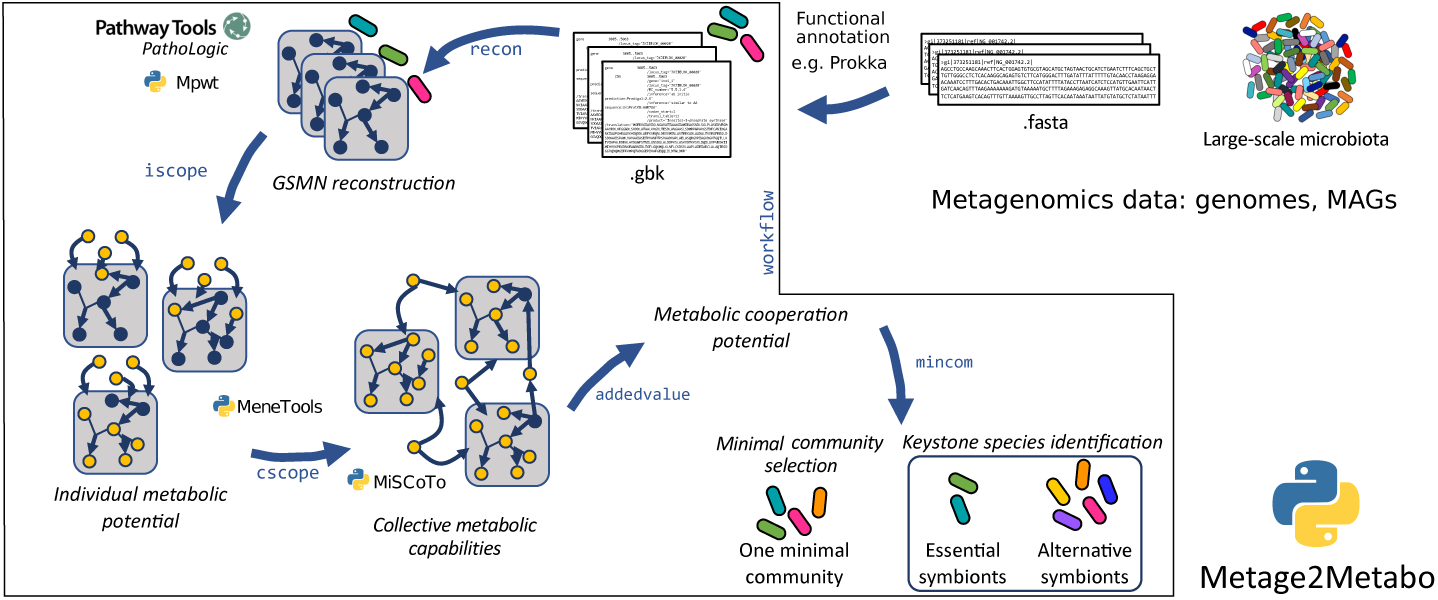
Overview of the M2M pipeline. The pipeline takes as inputs a set of annotated genomes that can be reference genomes of metagenomics-assembled genomes. Starting from these annotated genomes, Pathway Tools can be run in parallel for a large number of organisms. The resulting metabolic networks are analysed individually and collectively to identify metabolic capabilities, added-value of cooperation, and species of interest. The pipeline can be run as a whole or the steps can be performed individually.

The whole pipeline is called with the command *m2m workflow*. We detail below the characteristics of the M2M through a description of its main three steps.

### Large-scale metabolic network reconstruction

M2M enables a fast reconstruction of non-curated metabolic networks using Pathway Tools with the *m2m recon* command. It is the first multi-processing solution available to run this software and is therefore highly suitable to get metabolic insights into the hundreds or thousands of genomes that can be retrieved from metagenomic experiments.

Let us recall that Pathway Tools [27] is a graphical user interface (GUI) based software suite for the generation of GSMNs, called Pathway/Genome Databases (PGDBs). These PGDBs can be obtained from annotated genomes using Pathway Tools’s prediction component (PathoLogic) and curated afterwards. However, both Pathway-Tools GUI or command-line interface do not scale to the reconstruction of hundreds of GSMNs. With *m2m recon*, we propose an extension to Pathway Tools, that automatises the creation of these metabolic networks (in Systems Biology Markup Language (SBML) [22, 23] or PGDB the native Pathway Tools format) with PathoLogic, for large sets of genomes. Reconstructions are run in parallel, facilitating the scale-up of genome analysis.

To enable this reconstruction, we developed a Python wrapper named Mpwt (Multiprocessing Pathway Tools) that is included in M2M. This multiprocessing wrapper does not accelerate one PathoLogic run but it runs simultaneously multiple PathoLogic processes on different organisms. By default, Mpwt uses only one core but the user can allocate more to parallelise the runs. We recommend to give the number of available physical cores. Regarding memory requirements, they depend on the genome size but we advise to use at least 2 GB per core. The wrapper handles several types of genomic inputs (Genbank, Generic Feature Format (GFF) or PathoLogic format) and creates the input files needed by Pathway Tools. These files are then used by PathoLogic processes (one process by physical core) to create PGDBs. When all PGDBs are generated, Mpwt uses a lisp command of Pathway Tools to export the PGDB in an attribute-value flat files. These files are then used by the Padmet library [1] to generate SBML files suitable for exporting and sharing metabolic networks, for example for their use in other software for refinement.

### Analysis of metabolites producibility and potential added-value of cooperation

Individual metabolic network analysis is a first step to compare the quality of the reconstructions, as well as the functional potential of all species in a given environment. As metabolic networks are automatically reconstructed from thousands of possibly incomplete genomes, constraint-based methods relying on flux balance analysis [40] are not suitable. We therefore chose to use the network expansion algorithm [14] to assess the metabolic potential of each species. The network expansion algorithm computes the scope of a metabolic network from a description of the growth medium (seeds), that is, the family of metabolic compounds which are reachable according to a boolean abstraction of the network dynamics assuming that cycles cannot be self-activated. This algorithm has been widely used to analyse and refine metabolic networks[35, 31, 10, 45, 43], including for microbiota analysis [9, 38, 39, 18]. As the scope ignores the stoichiometry of metabolites involved in reactions, it appears to be a good trade-off between the accuracy of metabolic predictions and the precision required for the input data. It is therefore adapted to the difficulties met when studying the metabolism of hundreds or thousands of non-model organisms.

The analysis of metabolism provided by M2M can be called with the *m2m iscope* command. It predicts the set of reachable metabolites, the scope, in a metabolic network, starting from a set of seeds. This individual metabolic capability is calculated for each GSMN. All the scopes are exported as a json file and a summary is provided to the user: the intersection (metabolites reachable by all GSMN) and the union of all scopes, as well as the average size of the scopes, the minimal size and the maximal size of all, to get a glance at the range of metabolic capabilities among the species. This step is based on features implemented in Python packages that are also available as stand-alone tools: Menetools and MiSCoTo [1, 18].

In addition to individual studies, all metabolic networks are studied as a whole, to get insights into the complementarity of their metabolic pathways. Metabolic capabilities of the whole microbiota [18] can be computed using the network expansion algorithm with the command *m2m cscope*. This simulates the sharing of metabolic biosynthesis through a meta-organism composed of all GSMNs, and assesses the metabolic compounds that it can reach.

Based on the individual and community metabolic potentials, the global expected added-value (*m2m addedvalue*) of cooperation in a microbiota consists in the set of metabolites that can only be produced if several organisms share their metabolic biosynthesis. We advise the user to look at the list of producible compounds proposed by M2M, as it can be that some metabolites are false positive that do not necessitate cooperation for production, but were nonetheless selected due to missing annotations in the initial genomes. The *m2m addedvalue* command computes the identification of these compounds by comparing the results of individual and community scopes. It creates a target SBML file with these newly producible metabolites, for a possible use of these compounds in a community selection step.

### Identification of keystone species

This step, called with the *m2m mincom* command provides the reduction of the initial large-scale community through the identification of minimal size communities to ensure a metabolic objective is met. The latter is the reachability of metabolic compounds (targets) starting from nutrients (seeds) under the network expansion algorithm. The targets can be the added-value of cooperation as presented above, or subcategories of them as defined by the user. This step relies on a functionality of MiSCoTo [18] to propose a minimal community of organisms suitable for the objective. *M2m mincom* assumes that all metabolic transports have equal costs as transport reactions are not well identified in automatically reconstructed GSMNs. Further analysis introducing different costs based on additional knowledge on transport reactions can be performed by using the MiSCoTo package [18].

Many equivalent minimal communities are expected to exist but their enumeration can be computationally fastidious, as well as their analysis. An originality of M2M is to efficiently sample the space of solutions without the need for a full enumeration, thanks to the underlying logic programming solving. The intersection of solutions i.e. species occurring in every minimal communities or *essential symbionts* are computed. We describe as *keystone species* the organisms occurring in at least one minimal community (union of solutions). Keystone species therefore contain the essential and *alternative symbionts*, the latter occuring in some minimal communities but not all of them. These terms were inspired by the terminology used in flux variability analysis [40] for the description of reactions in all optimal flux distributions.

## Results

M2M was applied to two metagenomic experiments: a set of 1,520 bacterial high-quality draft reference genomes from the gut microbiota presented in [61], and 913 MAGs from the cow rumen published in [52]. The gut dataset consists in genomes of culturable bacteria, isolated from a large number of fecal samples and assembled into 338 species-level clusters. They cover major phyla: 796 Firmicutes, 447 Bacteroidetes, 235 Actinobacteria, 36 Proteobacteria and 6 Fusobacteria. We show that M2M is applicable to both types of datasets. We analyse in the next subsections the reconstructed GSMNs, the computation of the potential cooperation added-value, and keystone species. Finally we provide further analyses on the gut dataset with a deeper study of minimal communities composition.

The whole M2M workflow (from GSMN reconstruction to keystone species computation) took 155 minutes for the gut microbiota dataset. This reasonable time observed for computation is confirmed with the cow rumen dataset which ran in 81 minutes. In both cases, they were run on a cluster with 72 CPUs and 144 Go of memory.

### GSMN reconstruction

The genomes from the cow rumen were not annotated, a required feature for metabolic network reconstruction with M2M. We therefore annotated them using Prokka (v. 1.13.4) [47] as a preliminary step.

Results for the GSMNs reconstructions of both datasets and the analysis of individual metabolic potentials are presented in Table 1. The universe of metabolic reactions included in the reconstructions is of size 3,932 for the gut, and 4,418 for the rumen. Likewise, the universe of metabolites is of size 4,001 for the gut dataset, 4,466 for the rumen. The gut metabolic networks contained in average 1,144 (*±* 255) reactions and 1,366 (*±* 262) metabolites. 74.6% of the reactions were associated to genes, the remaining being spontaneous reactions or reactions added by the PathoLogic algorithm (they can be removed in M2M using the *–noorphan* option). GSMNs of the rumen dataset consisted in average of 1,155 (*±* 199) reactions and 1,422 (*±* 212) metabolites. 73.8% of the reactions were associated to genes. Supplementary Figure 1 displays the distributions of the numbers of reactions, pathways, metabolites and genes for both datasets. Altogether, these distributions are very similar for both datasets although the initial number of genes in the whole genomes varies a lot (Supp Fig. 1 g), a difference that is expected between MAGs and reference genomes. Interestingly, the average number of reactions per GSMN is slightly higher for the MAGs of the rumen than for the reference genomes of the gut. However, the smallest GSMN size is observed in the rumen (340 reactions vs 617 for the smallest GSMN of the gut). The similarity in the characteristics displayed by both datasets suggests a level of quality of the rumen MAGs close to the one of the gut reference genomes regarding the genes associated to metabolism.

**Table 1:** Results of the GSMN reconstruction step and metabolic potential analysis for two datasets (Avg = Average, “*±*” precedes standard deviation)

|  | Gut dataset | Rumen dataset |
| --- | --- | --- |
| initial data | draft reference genomes | MAGs |
| number of genomes | 1520 | 913 |
| **GSMN reconstruction** |  |  |
| all reactions | 3932 | 4418 |
| all metabolites | 4001 | 4466 |
| avg reactions per GSMN | 1,144 (± 255) | 1,155 (± 199) |
| avg metabolites per GSMN | 1,366 (± 262) | 1,422 (± 212) |
| avg genes per mn | 596 (± 150) | 543 (± 107) |
| % reactions associated to genes | 74.6 (± 2.17) | 73.8 (± 2.61) |
| avg pathways per mn | 163 (± 49) | 146 (± 32) |
| **metabolic potential** |  |  |
| number of seeds | 93 | 26 |
| avg scope per mn | 286 (± 70) | 101 (± 44) |
| union of individual scopes | 828 | 368 |

For both experiments, we designed a set of seeds metabolites representing a nutritional environment that is required for the metabolic analyses. It consists in components of a classical diet for the gut microbiota (93 metabolites), and basic nutrients (26 metabolites including inorganic compounds, carbon dioxide, glucose and cellobiose) for the rumen (Supp Tables 1 and 2). The scope represents the metabolic potential (reachable metabolites) starting from available seeds (nutrients) according to the network expansion algorithm [14], that is a simulation of a boolean abstraction of the GSMN dynamics. The average size of the individual scopes is relatively small compared to the universe of metabolites for both datasets, which is highly dependent to the chosen seeds and the potential gaps in the GSMNs. The union of all individual scopes is of size 828 and 368 for the gut and the rumen respectively (21 % of the universe of compounds for the gut, 8.2 % for the rumen). Supplementary Figure 1 (h, i, j and k) displays the distributions of the scopes for both datasets.

Altogether, these results suggest that despite a good size of GSMN reconstructions, the individual metabolic potential of each species is relatively small. This can be an impact of the choice of seeds nutrients, but it can also be due to missing annotations or gaps in the networks. In the latter case, it is likely that metabolic cooperation between the species fills the gaps of biosynthesis pathways and enables the putative producibility of more metabolites, which can be calculated with M2M.

### Added value of metabolic cooperation in the datasets

The metabolism of a given bacterium can be completed by the metabolism of others by filling gaps that exist in the first one, and therefore enabling the activation of more reactions than what can be expected when considered in isolation. By taking into account the complementarity between GSMNs in each dataset, it is possible to capture the benefit of metabolic cooperation over the producibility of metabolic compounds. Running *m2m cscope* evidenced that 296 and 156 new metabolites are potentially producible by the rumen GSMNs and the gut GSMNs respectively if cooperation between their members is allowed. This increases up to 15% and 25% the proportion of reachable metabolites in the whole universe of metabolites in the rumen and in the gut respectively. These are the metabolites that could not be reached by any GSMN when considered individually and therefore require shared metabolic capabilities to be produced.

We analysed the composition of the 156 newly producible metabolites for the gut dataset using the ontology provided for metabolic compounds in the MetaCyc database [26]. We divided the metabolites into 6 categories: amino acids and derivatives (5 metabolites), aromatic compounds (11), carboxy acids (14), coenzyme A (CoA) derivatives (10), lipids (28), sugar derivatives (58) (Supp. Table 3). The remaining 30 compounds were highly heterogeneous, we therefore restrained our subsequent analyses to subcategories of homogeneous targets. By default, the M2M pipeline sets all newly producible metabolites as targets to perform community reduction in the following steps. Yet, it is possible to change them by selecting a subset of these metabolites, for example the subcategories of targets as we will present in the next paragraphs.

### Community reduction

After the prediction of the cooperation added-value, M2M performs a community selection step based on a metabolic objective i.e. a list of metabolic compounds. It computes the size of a minimal community to ensure their producibility and their associated expected keystones species based on the GSMNs contents.

M2M proposes a community composition for the objective. Yet given the redundancy of functions in microbiotas [37], more than one minimal community solution is expected to exist. There might be thousands of them, and it can can be computationally difficult to enumerate due to the high combinatorics of the problem, especially for large sets of targets. Thanks to the logic programming solving assets of M2M, keystone species can be calculated without the need for all solutions to be enumerated, which is highly efficient computationally.

Applied to our datasets, *m2m mincom* led to a minimal community of size 25 to produce the 156 targets of the gut microbiota (Table 2). 11 members are essential symbionts, i.e. found in every minimal community, and the total number of keystone species is 205. Therefore, all minimal communities are composed of the same 11 members and a set of 14 others picked among 194 alternative symbionts. For the rumen dataset, the size of a minimal communities was 44, with an intersection of 20 essential symbionts and 107 alternative ones. The main pipeline of M2M stops after the computation of these sets of GSMNs, predicted to be relevant regarding the metabolic objective provided as input.

**Table 2:** Results of the community reduction step for the gut and rumen datasets. Keystone species occur in at least one of all equivalent minimal communities. Essential symbionts belong to all the minimal communities. Alternative sym<u>bionts belong to some but not all minimal communities</u>

|  | Gut dataset | Rumen dataset |
| --- | --- | --- |
| minimal size of community | 25 | 44 |
| keystone species | 205 | 127 |
| essential symbionts | 11 | 20 |
| alternative symbionts | 194 | 107 |

#### Keystone species in the gut dataset

We ran the community reduction step (*m2m mincom* command) with the 6 groups of targets that we isolated from the cooperation added-value for further analyses. The keystone species were computed for each group, and we studied their compositions in terms of phyla. The number of keystone species varies between 59 and 227, which is a strong reduction compared to the initial number of 1,520 GSMN used for the analysis.

#### Analysis of minimal communities in the gut

We enumerated all minimal communities for each individual group of targets using features of *m2m analysis*. The number of optimal solutions (size of the enumeration) is large, reaching more than 7 million equivalent minimal communities for the sugar-derivated targets (Table 3). We observe however that the size of the minimal community is quite small for each targets group (between 4 and 11).

**Table 3:**
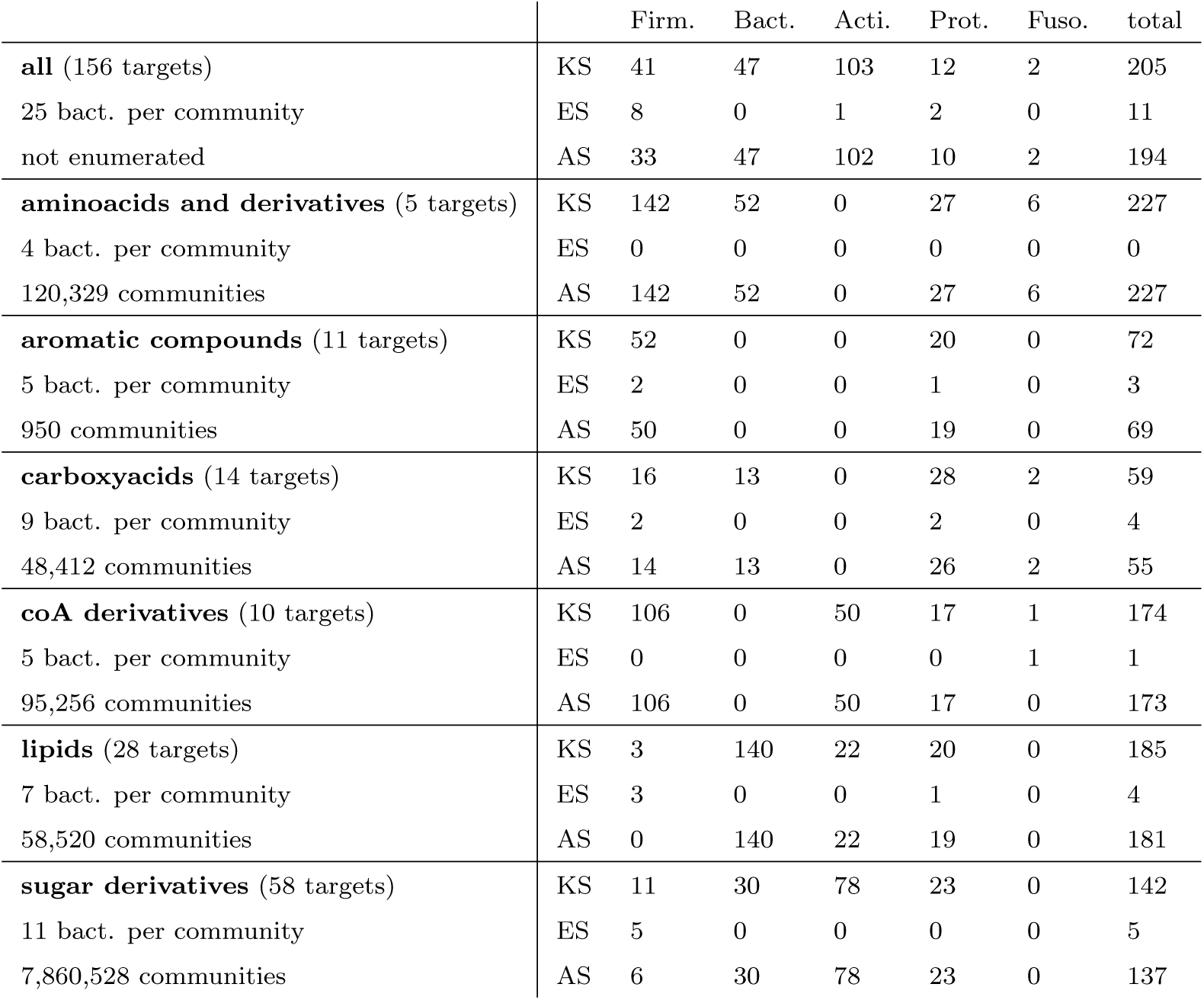
Community reduction analysis of the target categories in the gut. All minimal communities were enumerated, starting from the set of 1,520 GSMNs. KS: keystone species, ES: essential symbionts, AS: alternative symbionts, Firm.: Firmicutes, Bact.: Bacteroidetes, Acti.: Actinobacteria, Prot.: Proteobacteria, Fuso.: Fusobacteria.

The sizes of the enumerations make them difficult to analyse. Yet, our analyses suggest that the large number of optimal communities comes from numerous possibilities of combinatorial choices among a rather small family of bacteria. More precisely, the identification of keystone, essential and alternative groups of species to capture the diversity of the minimal communities yielded to the results depicted in Table 3. It gathers the composition of the three groups for the whole set of targets and the targets categories (see Supp. Tables 5 to 10 for the contents of the groups). In particular, essential symbionts are of high importance in minimal communities as they are found in each solution. More generally, compositions vary across the targets categories: a high proportion of keystone species for the production of lipids targets are Bacteroidetes whereas Firmicutes are highly represented for aminoacids and derivatives production.

Importantly, the keystones species for the categorical targets are not subsets of the ones for the whole set of targets. Large and heterogeneous sets of targets will lead to the selection of keystone species with more diversified biosynthesis pathways, likely to unblock the producibility of more compounds. On the contrary, for smaller and more homogeneous sets of targets, it is likely that a larger number of organisms with equivalent biosynthesis capabilities will be selected. Indeed, the number of keystone species is not necessarily smaller for small groups of targets.

#### Evidencing organisms with equivalent roles in microbial communities

In order to visualise the association of GSMNs in individual solutions, we created a graph whose nodes are the keystone species, and whose edges represent the association between two members if they co-occur in at least one of the enumerated communities. We created such graphs for each of the targets sets (lipids, sugar derivatives, aromatic compounds, aminoacids derivatives, carboxyacids and coA derivatives). Yet, they were very dense (185 nodes, 6888 edges for the lipids, 142 nodes and 6602 edges for the sugar derivatives), which is expected given the large number of optimal communities and the relatively small number of keystone species.

We compressed these graphs into power graphs to capture the mechanisms underlying the combinatorics of associations within minimal communities. Power graphs enable a lossless compression of re-occurring motifs within a graph: cliques, bicliques and star patterns [44]. We generated them using PowerGrASP [6] and visualised them with Cytoscape (version 2.8.3) [49] and the CyOog plugin (version 2.8.2) developed by [44]. As compressed graphs have a better readability, the power graphs generated from the association graphs of the enumerations enable to pinpoint metabolic equivalency between members of the keystone species.

Figure 2 presents the compressed graphs for each set of targets. Graph nodes are the keystone species, coloured by their phylum. A version of the figures with nodes identification is available in Supplementary Figures 2 to 7 (see. Supp. Table 4 for a mapping between identifiers and taxonomy). Nodes are included into power nodes, connected by power edges, depicting the equivalency between species with respect to the enumerated solutions. Symbionts belonging to a power node play the same role in the construction of the minimal communities. Essential symbionts are conveniently represented in the power graphs, either into power nodes with loops (Fig 2 a, e) or individual nodes connected to power nodes (Fig 2 a, c, d, f).

**Figure 2:**
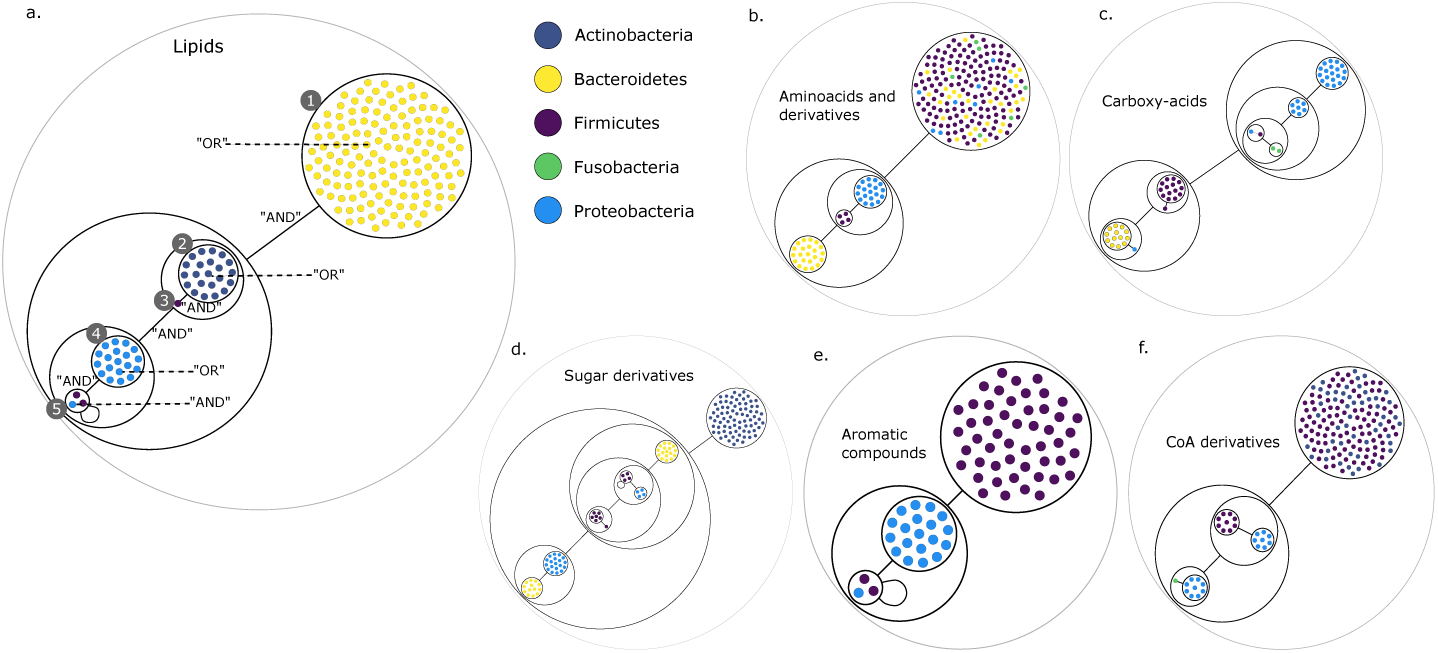
Network analysis of microbial associations within communities for the gut dataset. Each category of metabolites predicted as newly producible in the gut was defined as a target set for community selection among the 1,520 GSMNs from the gut dataset. For each metabolic group, keystone species and the full enumeration of all minimal communities were computed. Association graphs were built to associate members that are found together in at least one minimal community among the enumeration. These graphs were compressed as power graphs to identify patterns of associations. Power graphs a., b., c., d., e., f., g. present the patterns of associations for lipids, aminoacids and derivatives, carboxy-acids, sugar derivatives, aromatic compounds, and coenzyme A derivative compounds respectively. Nodes colours describe the phylum the initial genomes belong to. Figure a. has an additional description to ease readability. Edges symbolise conjunctions (”AND”), the co-occurrences of nodes in regular powernodes (as in powernode 1, 2, 4) symbolise disjunctions (”OR”) related to alternative symbionts. Powernodes with a loop (e.g. powernode 5) indicate conjunctions. Therefore, each enumerated minimal community for lipids production is composed of the two Firmicutes and the Proteobacteria from powernode 5, the Firmicutes node 3 (the four of them being the essential symbionts), and one Proteobacteria from powernode 4, one Actinobacteria from powernode 2 and 1 Bacteroidetes from powernode 1.

We observe that power nodes often contain GSMNs from the same phylum, indicating that an interesting function is shared by the taxonomic group or possibly the species-level clusters of genomes [61], for the producibility of the targets. The power graph gives information on the assembly of the GSMNs in these solutions. The power graphs for the groups of targets presented here are particularly informative to capture the composition of communities. Figure 2 a has additional comments to ease the reading. Each minimal community is composed of one Bacteroidetes from power node (PN) 1, one Actinobacteria from PN 2, the Firmicute member 3, one Proteobacteria from PN 4 and finally the two Firmicutes and the Proteobacteria from PN 5. For all the targets groups of this study, the numerous enumerations can be summarised with a boolean formula derivated from the graph compressions. For instance for the lipids of Figure 2 a, the community composition is the following (∨*PN* 1) ∧ (∨*PN* 2) ∧ (*PN* 3) ∧ (∨*PN* 4) ∧ (∧*PN* 5). This visualisation of community compositions thus enables a better understanding of the associations of organisms into the proposed communities, and the relevance of keystone species to identify members with functions of interest for the producibility of targets.

## Discussion

In this paper we present a new software for the functional analysis of metagenomic datasets at the metabolic level. M2M reconstructs metabolic networks using the Pathway Tools software and an efficient multi-processing wrapping. These GSMNs are then analysed individually and collectively to compute the potential added-value of metabolic cooperation and its associated minimal communities. Thanks to a graph-based modeling of producibility and powerful logic programming solving approaches, the whole space of minimal communities solutions can be parsed to retrieve all interesting bacteria, that we call keystone species, with respect to the metabolic objective. M2M can therefore suggest species for further analyses such as targeted curation of metabolic networks and deeper analysis of the genomes, that cannot be contemplated in an automatic way for the hundreds or thousands genomes of a microbiota.

The functionality of metagenomic sequences can be analysed at higher levels by directly computing functional profiles from reads [17, 51, 50, 42]. However, metabolic network reconstruction provides a more thorough study of the species by gathering the information about the reactions and pathways (complete or incomplete) they catalyse. Thanks to the improvements of the software suites for GSMN reconstruction, the automatically-built drafts provide a detailed view of the metabolic capabilities of species even without manual curation, although the latter is still needed for further analyses and accurate quantitative simulations of the metabolism.

Here we chose the Pathway Tools suite and its PathoLogic software [27] for GSMN reconstruction. It is based on the MetaCyc database [7] that covers a large taxonomic range of organisms and groups reactions into pathways that are convenient for a higher-level analysis of metabolism. Pathway Tools is distributed with a GUI that is very user-friendly for manual curation and analysis of a small number of GSMNs, but is limiting when aiming at reconstructing a large number of them. The multi-process wrapping we propose in this paper is entirely command-line based and performs metabolic network reconstruction for a large number of species in a transparent way for the user. The resulting GSMNs can also be opened and refined with Pathway Tools GUI.

The assets of Pathway Tools are the decomposition of GSMN into pathways and the possibility to only use PathoLogic without further automatic gap-filling refinements to the model. The latter can be non-indicated since the species under study are not expected to be self sufficient for growth in a microbiota environment. Therefore we advocate for the use of GSMN drafts that can be likely to miss metabolic reactions, but have a lower risk of false positive reactions. The latter would otherwise possibly be added by automatic curation and gap-filling to sustain individual growth of the models. False positive reactions may lead to missed interactions between species when considering metabolic cooperation. GSMN of interest after M2M analysis can be further analysed and refined if needed. Nevertheless, there are several other software available for GSMN reconstruction in addition to Pathway Tools such as Kbase [3], ModelSEED [21] or CarveMe [32]. Metabolic networks resulting from such platforms can be used as inputs to M2M for metabolic analysis as the reconstruction step can be bypassed and the tool accepts GSMNs in SBML format [23], widely used for exchanging and distributing models.

Functional analysis of metabolic networks using the qualitative criterion of the scope has been demonstrated to be robust and relevant [20, 10, 43]. The scope provides a snapshot of producible metabolites, thus identifying the sub-network that can be activated under given nutrient conditions. It is computed with the network expansion algorithm. A main difficulty can be to build the set of seeds that are available as nutrients for the analysis and on which the computation of network expansion relies. In particular, the algorithm has been demonstrated to be sensitive to cycles in GSMNs and it is therefore relevant to include some cofactors (e.g. ATP) in the seeds to activate such cycles, the way many studies proceed [12, 19, 15, 25]. Network expansion is a good trade-off with respect to quantitative constraint-based methods such as flux balance analysis [40] as it does not require biomass reactions nor accurate stoichiometry, and it can easily scale to thousands of networks considered in interaction. The cost of exchanges is not taken into account in the M2M pipeline as transport reactions are hardly recovered by automatic methods [5]. Yet, the standalone MiSCoTo package used in M2M has an option for taking into account exchanges. It can compute communities while minimising the cost of exchanges, and suggest them although it comes with a computational cost.

The number of curated metabolic networks for species found in microbiotas is in growing evolution [33]. They are a highly valuable resource for the study of interactions between members of communities. Yet the variety of (reference) genomes obtained from shotgun metagenomic experiments is such than species and strains vary a lot and may not belong to the ones for which a curated GSMN is available. In addition, the rise of methods for assembling genomes directly from metagenomes (MAGs) leads to incomplete genomes for possibly unknown species on which one may still want to get metabolic insights [2]. Therefore, building de novo automatic metabolic network drafts for each genome or MAG is a relevant solution to globally analyse the functions of a metagenome, especially in the rapidly evolving context of the gut microbiota [59].

A critical issue in the identification of minimal communities is the large number of equivalent solutions and the fact that many species can play an equivalent role in these communities. Three strategies can be investigated. Targets can be divided into families to better understand the role of species with respect to their production. This can be achieved by relying on ontologies to classify metabolites. A second strategy is to identify keystone species and carefully distinguish essential and alternative symbionts. Finally, we presented an original method to visualise and summarise the large number of minimal communities given a metabolic objective by building power graphs [44]. It evidenced a taxonomic homogeneity in the clusters of species identified for groups of targets in the gut microbiota dataset. Altogether, Studying the metabolic potential of large communities is an iterative process that still requires biological expertise.

## Conclusions

In this paper we present a new software for the functional analysis of genomes and MAGs from metagenomic experiments by reconstructing and comparing metabolic networks. M2M automatically builds GSMNs starting from annotated genomes. The pipeline is scalable and efficient by performing the reconstruction in a multiprocess framework. The resulting metabolic networks are analysed to capture the set of metabolites they are expected to reach, either individually or collectively through metabolic cooperation. The added-value of cooperation in terms of increase of the producible metabolites set is computed, and minimal communities for this objective are calculated. The large combinatorics of minimal communities due to functional redundancy in microbiotas is addressed by providing the alternative and essential symbionts of all solutions without the need for a complete enumeration. The groups of species identified are relevant for deeper analyses and their size makes the following studies more tractable than when considering the whole dataset of initial genomes.

M2M can be used as a pipeline but each step can also be performed individually thus increasing the flexibility of analysis and the range of its applications. Our method is robust against the uncertainty inherent to metagenomics data and is, to our knowledge, the only one available to predict keystone species at the metabolic level starting from large sets of genomes that can originate from poorly-studied or unknown species. M2M can be contemplated for a use together with abundance data to understand the importance of the identified keystone species. It could also serve as a basis to understand the evolution of microbial composition of patients in longitudinal studies at the metabolic level. We believe this software is of interest as the number of available genomes from metagenomic studies continues to rise, entailing a need for scalable predictive methods that tolerate the incompleteness of data.

## Availability and requirements

**Project name**: metage2metabo

**Project home page**: https://github.com/AuReMe/metage2metabo

**Operating system(s)**: Linux and MacOS

**Programming language**: Python 3.6 or higher

**License**: GNU General Public License v3.0

**Any restrictions to use by non-academics**: Licence needed for the use of Pathway Tools

## List of abbreviations

M2M: metage2metabo
GSMN: Genome-Scale Metabolic Network
MAG: Metagenome-Assembled Genome
GFF: Generic Feature Format
RAM: Random-Access Memory
PGDB: Pathway Genome DataBase
SBML: Systems Biology Markup Language
GUI: Graphical User Interface
PN: Power Node

## Availability of data and material

The rumen dataset MAGs used for the experiments were downloaded from https://www.ncbi.nlm.nih.gov/assembly/?term=PRJEB21624. The gut microbiota genomes were downloaded from https://www.ncbi.nlm.nih.gov/assembly/?term=PRJNA482748

## Author’s contributions

ABe and CF developed the pipeline, designed and ran the experiments, performed the analyses. CF wrote the manuscript. MA and ABr designed technical solutions. AS supervised the work. All authors read and reviewed the manuscript.

## Acknowledgements

The authors acknowledge the GenOuest bioinformatics core facility for providing the computing infrastructure. We also acknowledge P. Karp, S. Paley, M. Krummenacker, R. Billington, A. Kothari from the Bioinformatics Research Group of SRI International for their help regarding Pathway Tools. Finally, we thank Lucas Bourneuf for his help on power graph analyses.

## Additional Files

**SuppFigures.pdf — Additional figures**

**SuppTables.xlsx — Additional tables**

