## Supplementary figures for "Metage2Metabo: metabolic complementarity applied to genomes of large-scale microbiotas for the identification of keystone species"

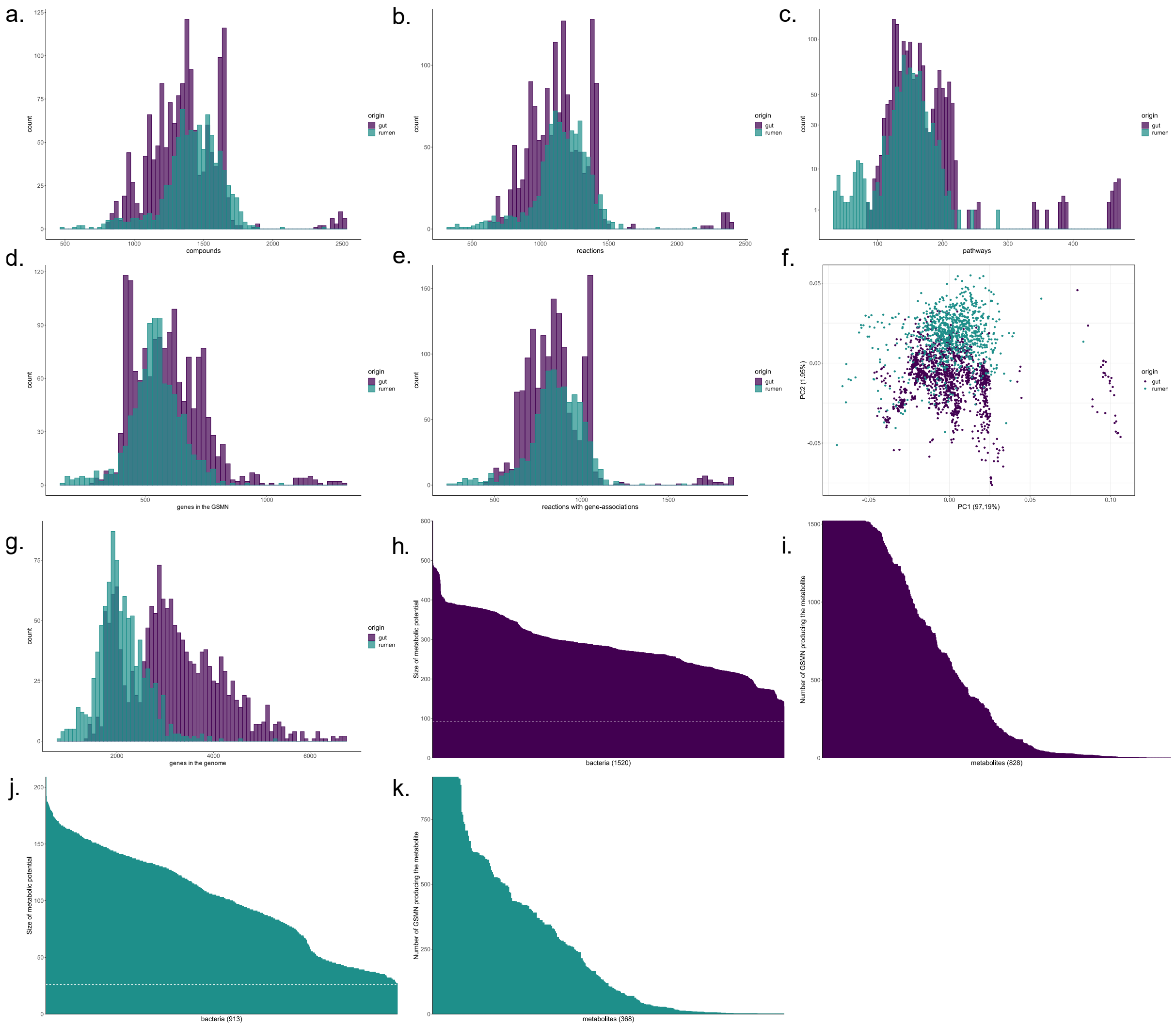


*Figure 1. Characteristics of the metabolic networks built for the gut and the rumen datasets.* ***a.*** *Distribution of the number of metabolic compounds in GSMNs reconstructed for the gut dataset (purple) and the rumen dataset (blue).* ***b.*** *Distribution of the number of metabolic reactions.* ***c.*** *Distribution of the number of complete pathways according to the MetaCyc database.* ***d.*** *Distribution of the number of genes included into the GSMNs.* ***e.*** *Distribution of the number of reactions associated to genes.* ***f.*** *Principal component analysis of the GSMNs reconstructions based on the previous characteristics (a. to e).* ***g.*** *Distribution of the number of genes in the initial genomes/MAGs.* ***h.*** *Individual metabolic potentials (scopes) for the gut bacteria, dotted line represents the number of seeds (nutrients) used in the algorithm.* ***i.*** *Reachability of metabolites by gut bacteria.* ***j.*** *Individual metabolic potentials (scopes) for the rumen bacteria, dotted line represents the number of seeds (nutrients) used in the algorithm.* ***k.*** *Reachability of metabolites by rumen bacteria.*

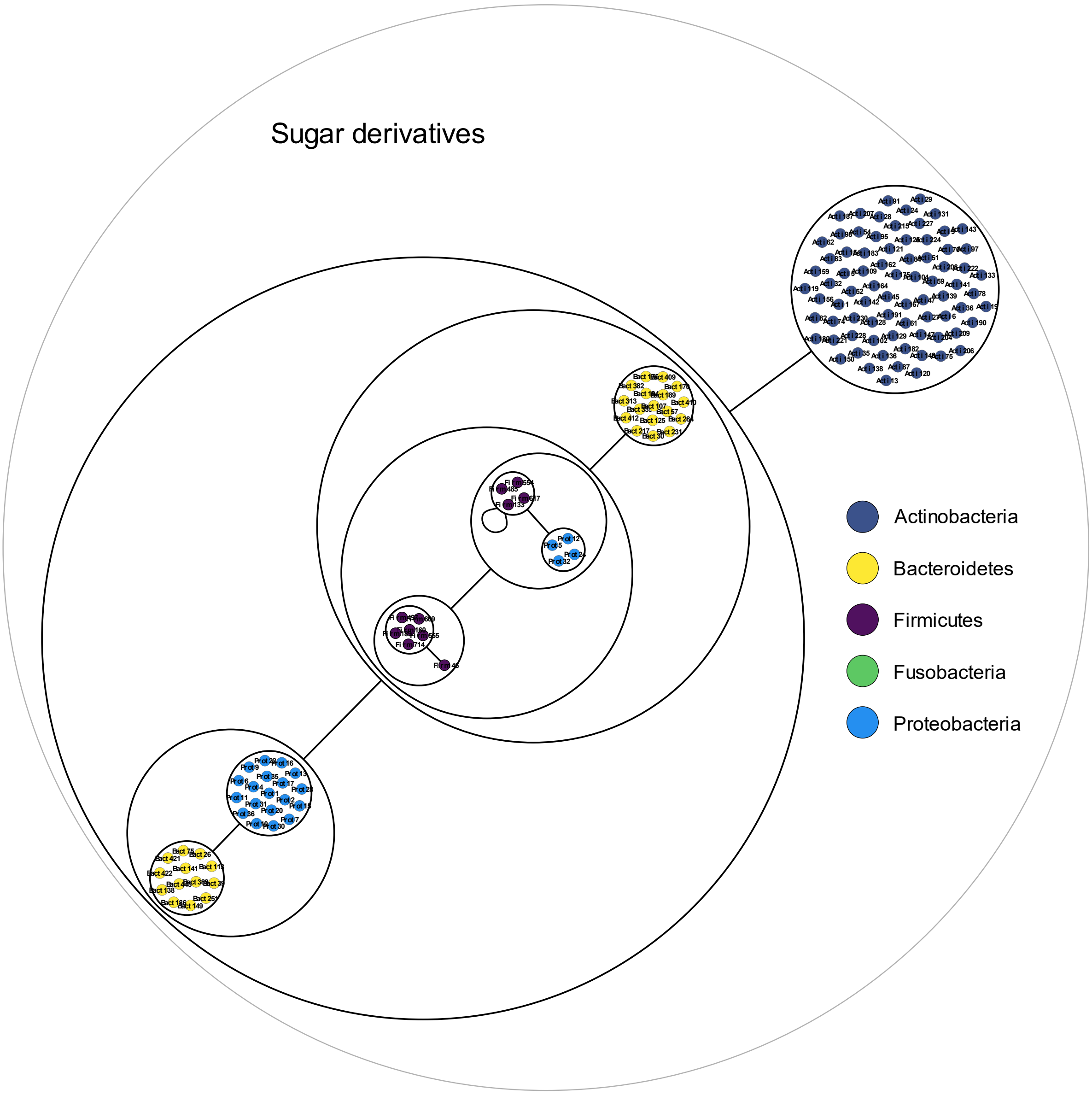


*Figure 2. Powergraph associated to the minimal communities producing the sugar derivatives group of targets*

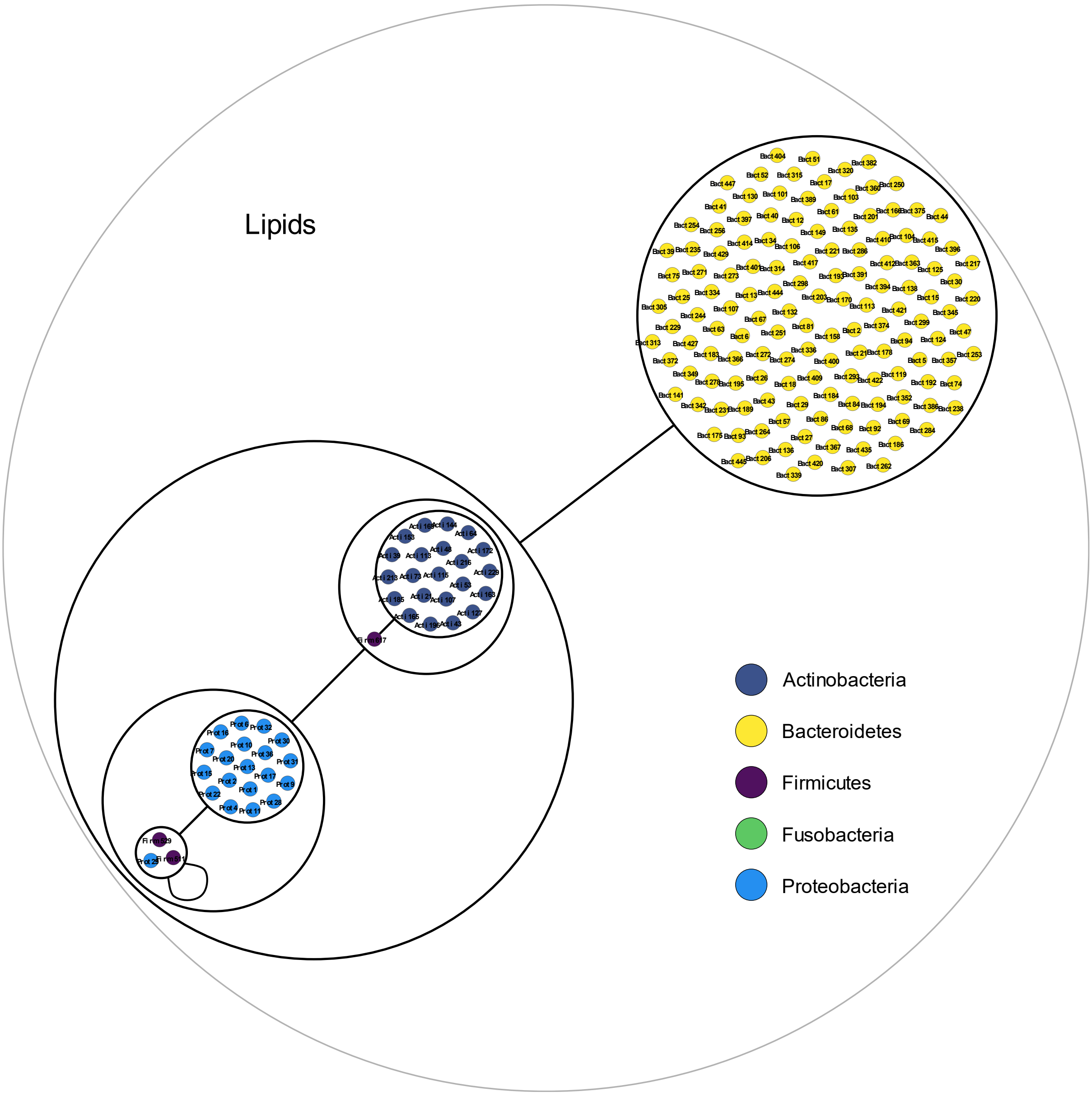


*Figure 3. Powergraph associated to the minimal communities producing the lipids group of targets*

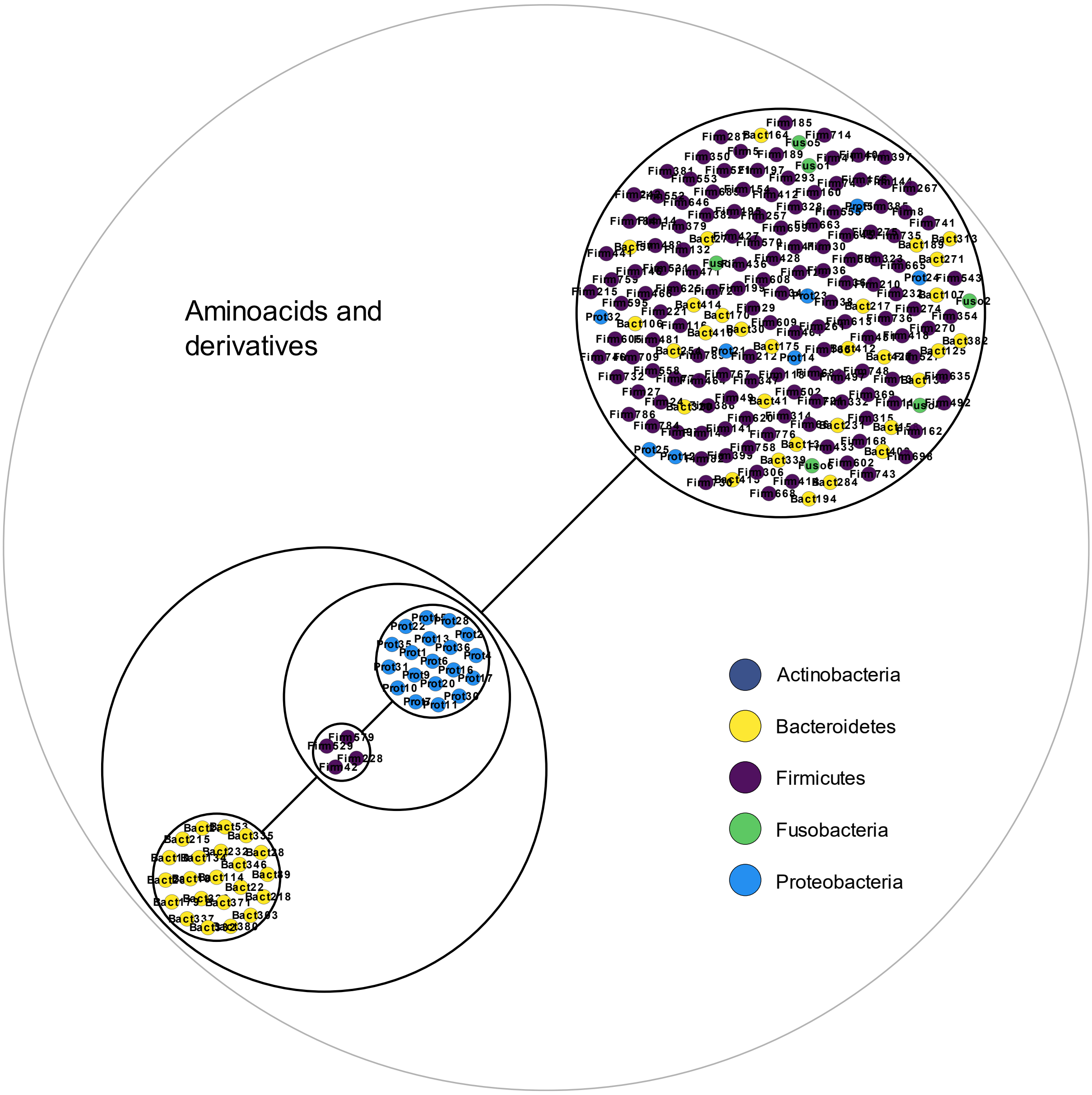


*Figure 4. Powergraph associated to the minimal communities producing the aminoacids and derivatives group of targets*

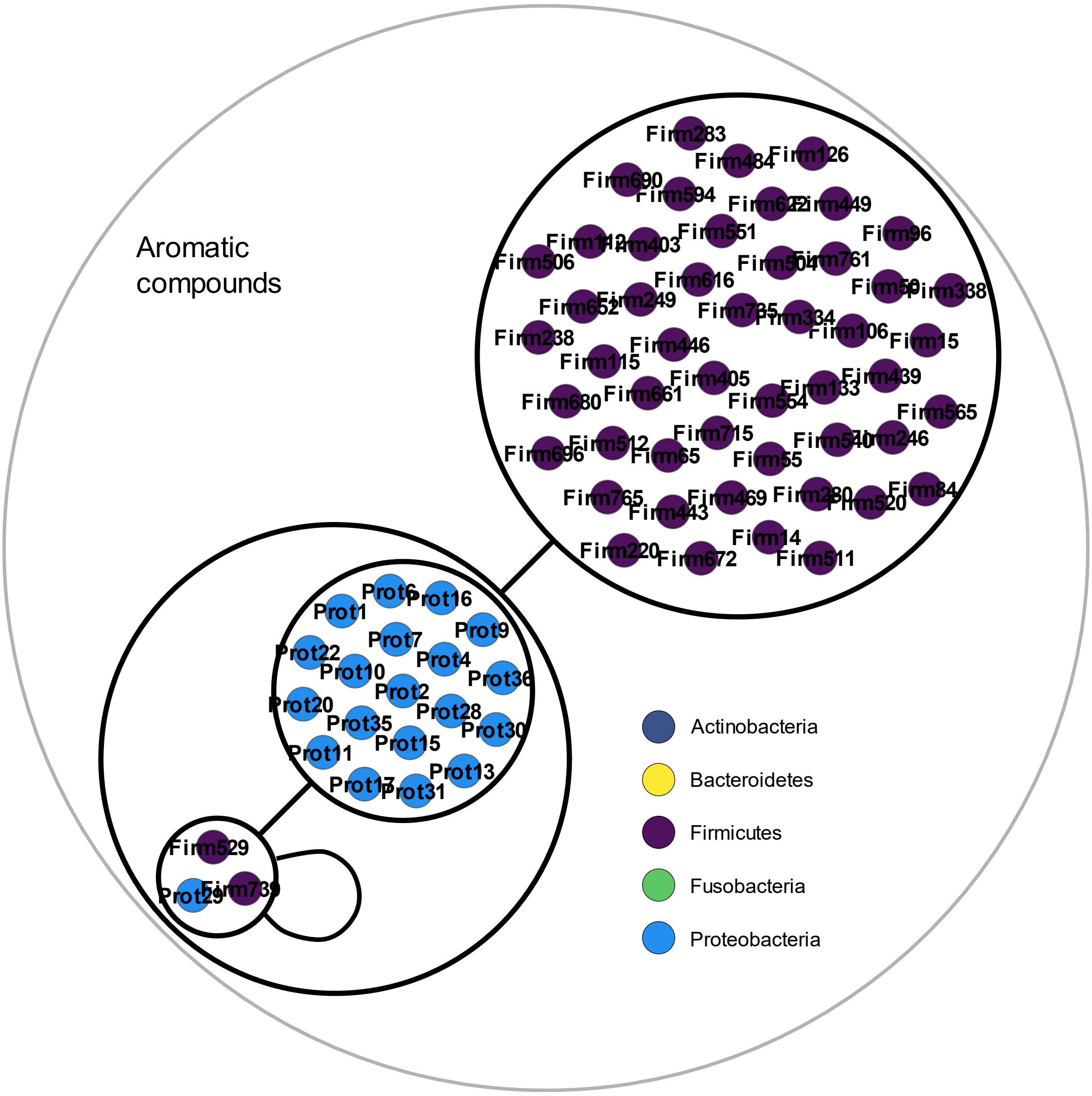


*Figure 5. Powergraph associated to the minimal communities producing the aromatic compounds group of targets*

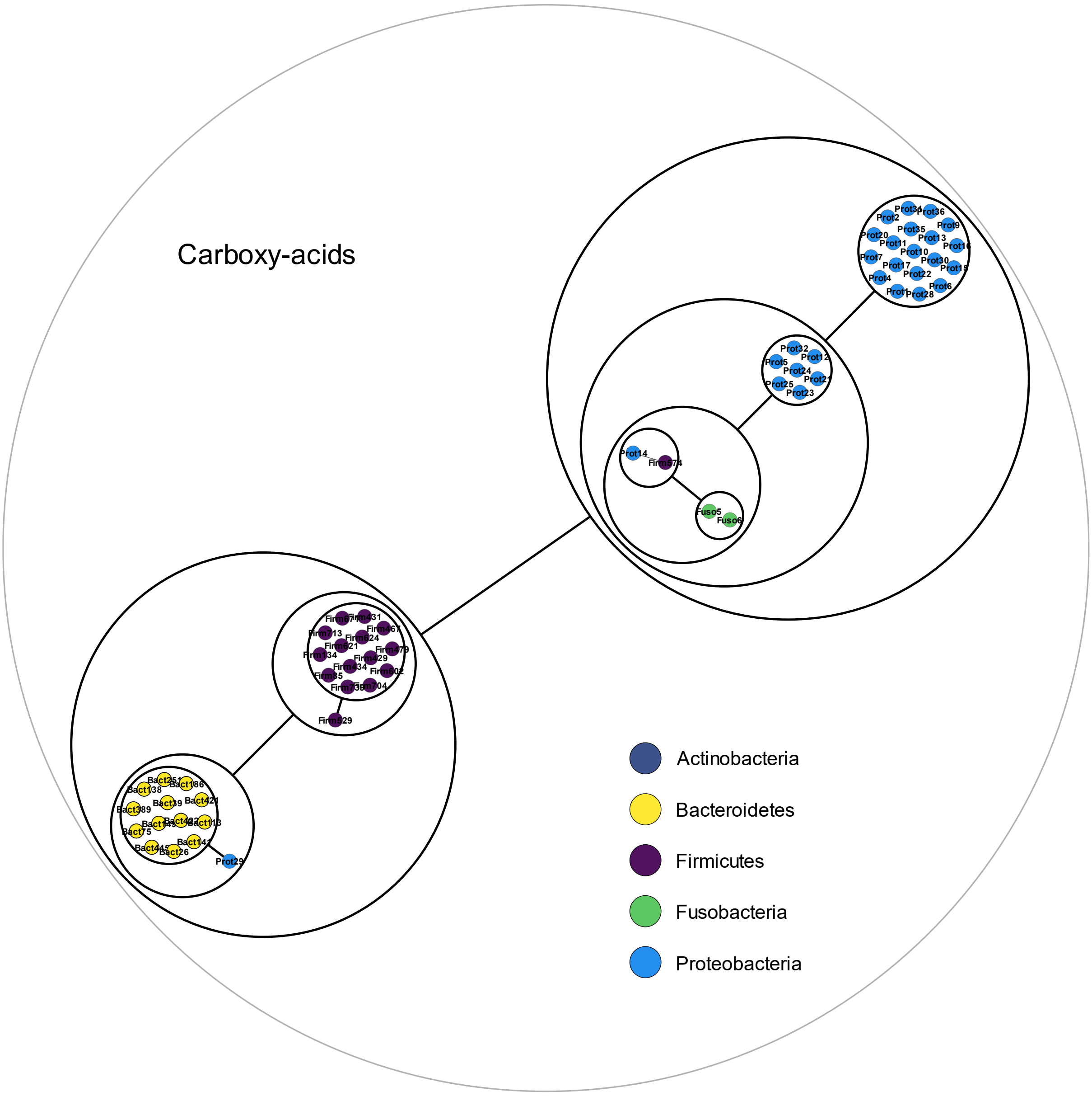


*Figure 6. Powergraph associated to the minimal communities producing the carboxy-acids group of targets*

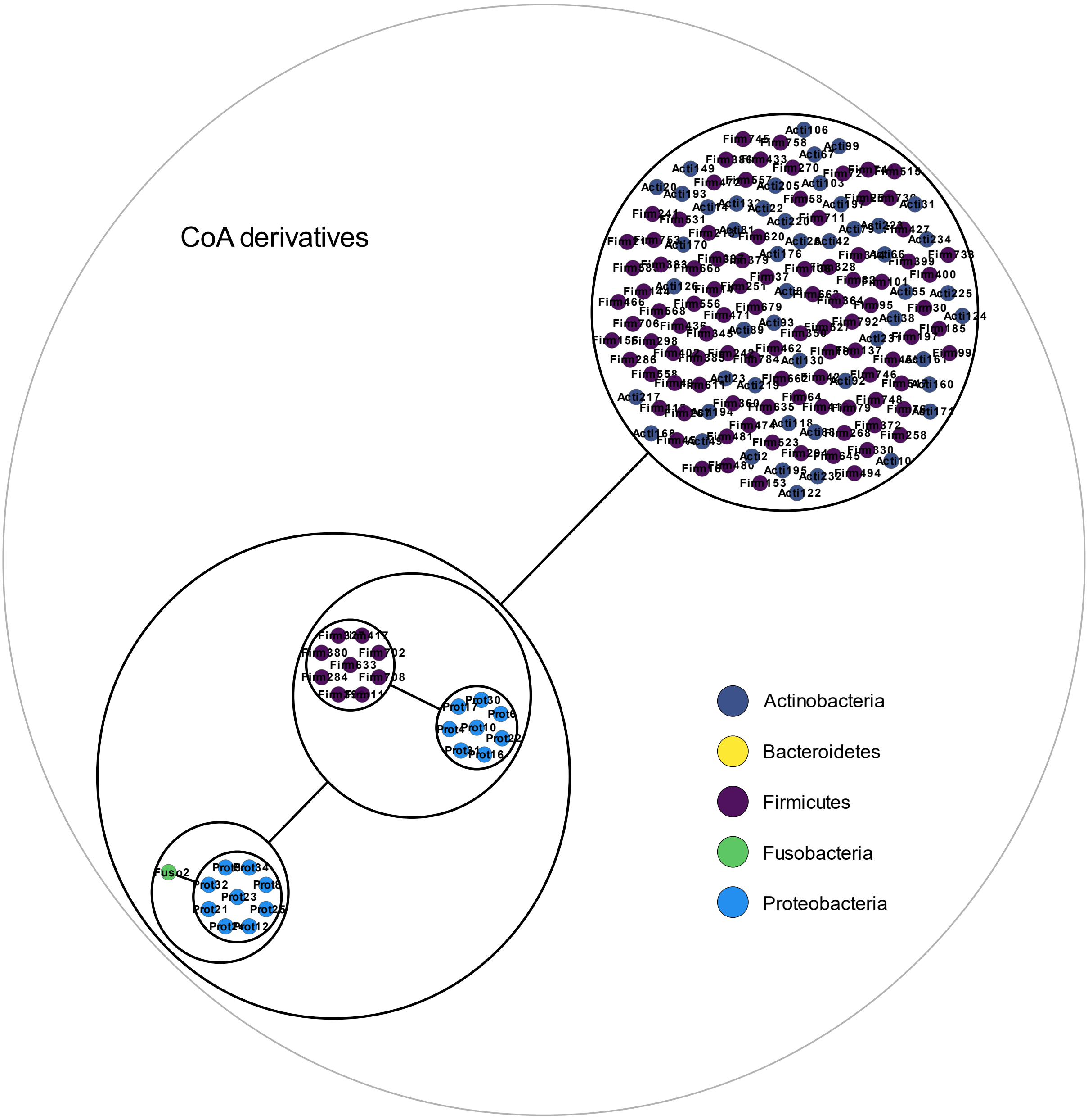


*Figure 7. Powergraph associated to the minimal communities producing the coenzyme A derivatives group of targets*
